# Nuclear aging in polyglutamine-induced neurodegeneration

**DOI:** 10.1101/2023.06.09.544056

**Authors:** Dina Pereira, Janete Cunha-Santos, Ana Vasconcelos-Ferreira, Joana Duarte-Neves, Isabel Onofre, Vítor Carmona, Célia A Aveleira, Sara M Lopes, Diana D Lobo, Inês M Martins, Nélio Gonçalves, Cláudia Cavadas, Luís Pereira de Almeida

## Abstract

Machado-Joseph disease (MJD) is an autosomal dominantly-inherited neurodegenerative disorder characterized by an over-repetition of the CAG trinucleotide of the *ATXN3* gene, conferring a toxic gain-of-function to the resulting ataxin-3 protein. Despite the significant advances produced over the last years, the molecular mechanisms involved in MJD are still unclear and no treatment able to modify the disease progression is available. Aging is the major risk factor for neurodegenerative disorders, being associated with the occurrence and progression of several diseases, such as Alzheimer’s, Huntington’s, among others. The nuclear membrane proteins - lamins - and lamin-processing related proteins, such as ZMPSTE24, have been shown to be altered, not only during normal aging, but also in neurodegenerative disorders, such as Alzheimer’s disease.

Taking this into account, we aimed at investigating the role of aging in MJD by evaluating the presence of age-related markers in human and animal MJD models. Decreased levels of lamins B and C, together with decreased ZMPSTE24 levels were identified in the different MJD models. Accordingly, abnormalities in nuclear circularity, a hallmark of aging, were also observed in a N2a MJD cellular model, supporting an age-related phenotype. Furthermore, overexpressing progerin, the abnormal lamin A, generated in Hutchinson Guilford Progeria Syndrome patients that present premature and accelerated aging, in a relevant brain area of a lentiviral MJD mouse model, induced an aggravation of MJD-associated neuropathology.

Our results suggest that aging is a key player in the context of MJD pathogenesis, unveiling new pathways for the development of future therapies for the disease.

## 1 INTRODUCTION

Aging can be defined as a natural time-dependent process, characterized by the progressive decline of biological functions, after the organism has achieved its maximal reproductive capacity, towards mortality (Farooqui & Farooqui, 2009). It is the major risk factor for neurodegenerative disorders, such as Alzheimer’s disease and other dementias, Parkinson’s disease, among others. Such disorders have caught special attention due to their irreversibility and absence of an effective treatment (reviewed in (Jin & Cai, 2023). Machado-Joseph disease or spinocerebellar ataxia type 3 (MJD/SCA3) is the most commonly inherited autosomal-dominant neurodegenerative disorder worldwide, caused by the pathological expansion of the CAG trinucleotide repeat in exon 10 of the *ATXN3* gene localized in the long arm of chromosome 14 (14q32.12) (Kawaguchi et al., 1994; Takiyama et al., 1993). As observed for the other eight, so far identified, members of the polyglutamine diseases, expression of the polyglutamine-expanded protein, mutant ataxin-3 in SCA3 case, leads to the accumulation of mutant protein inclusions and severe neuronal loss in several brain regions of the spinocerebellar tract and other brain areas, including the cerebellum, brainstem and basal ganglia (Alves et al., 2008; Rüb et al., 2008).

Although the genetic cause of MJD is well defined and described, and regardless of significant research progress that we and others provided during the last years, the mechanisms underlying neuronal degeneration are still largely unknown, and no treatment able to modify the disease progression is available (Duarte Lobo et al., 2020; Nóbrega et al., 2018; Vasconcelos-Ferreira et al., 2022). Nevertheless, a common feature among MJD and other polyglutamine diseases is the late onset of their clinical manifestation, which is still not fully understood (Mattson & Magnus, 2006). Aging itself is considered to be the principal risk factor for the onset of neurodegenerative disorders associated with proteostasis impairments and aggregate deposition (Knowles et al., 2014). Given the transversal characteristics of polyglutamine diseases, such as MJD, we speculate that aging might be implicated in these diseases, particularly at the level of the nuclear envelope integrity.

Prelamin A, a non-processed form of lamin A (one of the “building blocks” of the inner nuclear membrane envelope) was found to be accumulated in vascular smooth muscle of aged, but not in young individuals. This results from a decrease in the expression of ZMPSTE24 (involved in the maturation of lamin A), suggesting an age-dependent decline that gives rise to a phenotype similar to that seen in accelerated-aging syndromes such as Hutchinson-Gilford Progeria Syndrome (HGPS) (Ragnauth et al., 2010).

HGPS is a rare genetic condition in which children appear to age prematurely (Gilford, 1904; Hutchinson, 1886). It is classically caused by a *de novo* heterozygous mutation (1824 C>T, pG608G) in exon 11 of *LMNA* gene, which activates a cryptic splice donor site, leading to the production of a prelamin A mRNA with an internal deletion of 150 base pairs. This altered transcript is translated into a mutant lamin A protein, called progerin, that lacks 50 amino acids within its C-terminus including the FACE1/ZMPSTE24 cleavage site. Consequently, progerin remains permanently farnesylated and accumulates in an improper fashion at the inner nuclear membrane, inducing numerous cellular defects that are also considered as hallmarks of normal aging (Carrero et al., 2016; Eriksson et al., 2003). These features include highly blebbed nuclei with thickened lamina, accumulation of DNA damage and defects in telomere function (leading to differentiation defects and premature cellular senescence) among others (Carrero et al., 2016).

A link between laminopathies (disorders caused by defects in the nuclear lamina) or nuclear dispersion and age-related neurodegenerative disorders, namely Alzheimer’s disease, was also found (Chang et al., 2011; Frost, 2016). Moreover, neurons of tau transgenic Drosophila and of *postmortem* human brain tissue from Alzheimer’s disease patients have been shown to harbour significant nuclear invaginations and reduced lamin-B protein levels (another component of the nuclear lamina) when compared to controls (Frost, 2016).

In this work, we investigated the presence of age-related markers associated with nuclear integrity in cellular (patient fibroblasts and N2a cells) and animal models (transgenic and lentiviral mouse models) of MJD.

Study of human fibroblasts and from different genetic models of SC3/MJD revealed an age-related phenotype with decreased nuclear protein levels of lamins B and C, decreased ZMSPSTE24 mRNA and protein levels and abnormalities in the nuclear circularity, all “hallmarks” of aging. Importantly, overexpression of progerin aggravated MJD neuropathology, and associated neuroinflammation, suggesting that aging may have a significant role in MJD pathogenesis, aggravating MJD-associated neuropathology.

## 2 RESULTS

### 2.1 ZMPSTE24 downregulation in human and rodent MJD models

ZMPSTE24, a zinc metalloproteinase responsible for the cleavage of the farnesyl group in the last processing step of lamin A maturation, has been reported to be decreased in vascular smooth muscle cells and fibroblasts of aged individuals and centenarians, respectively, leading to the accumulation of prelamin A, and to a phenotype similar to that seen in HGPS (Lattanzi et al., 2014; Ragnauth et al., 2010).

The evidence of an age-related decline of this processing protein led us to investigate ZMPSTE24 mRNA and protein levels in three models of MJD: MJD human fibroblasts, the cerebellum of a MJD transgenic mouse model (Torashima et al., 2008) and the striatum of a MJD LV-based mouse model (Alves et al., 2008; de Almeida et al., 2002). Comparisons were performed with healthy counterparts.

As shown in Figure 1a, b and c, both mRNA and protein levels of ZMPSTE24 were significantly decreased (24.1± 8.7% and 64.2 ±16.4%, respectively) in fibroblasts from MJD patients, compared with fibroblasts from control individuals. MJD transgenic mice also revealed decreased protein levels of ZMPSTE24 with a significant alteration (47.2±16.5%) in mice with more than one year of age that was not observed in WT mice (Figure 1d and h), with no alterations seen at other time points in this mouse model (Figure 1e, f, g). In the striatum brain region, isolated from the MJD LV-based mouse model (Figure 1i), it was also possible to observe a significant decrease in *Zmpste24* mRNA levels at 4 weeks post-injection (30.5±8.4 %) (Figure 1j), and in the protein levels of mice, 8 weeks after lentiviral transduction (22.64 ± 7.2%) (Figure 1m). No alterations were found in the protein levels of mice with 4 weeks (Figure 1k) and mRNA levels in mice with 8 weeks of age (Figure 1l). Overall, these results suggest that ZMPSTE24 expression is compromised in MJD.

**Figure 1.**
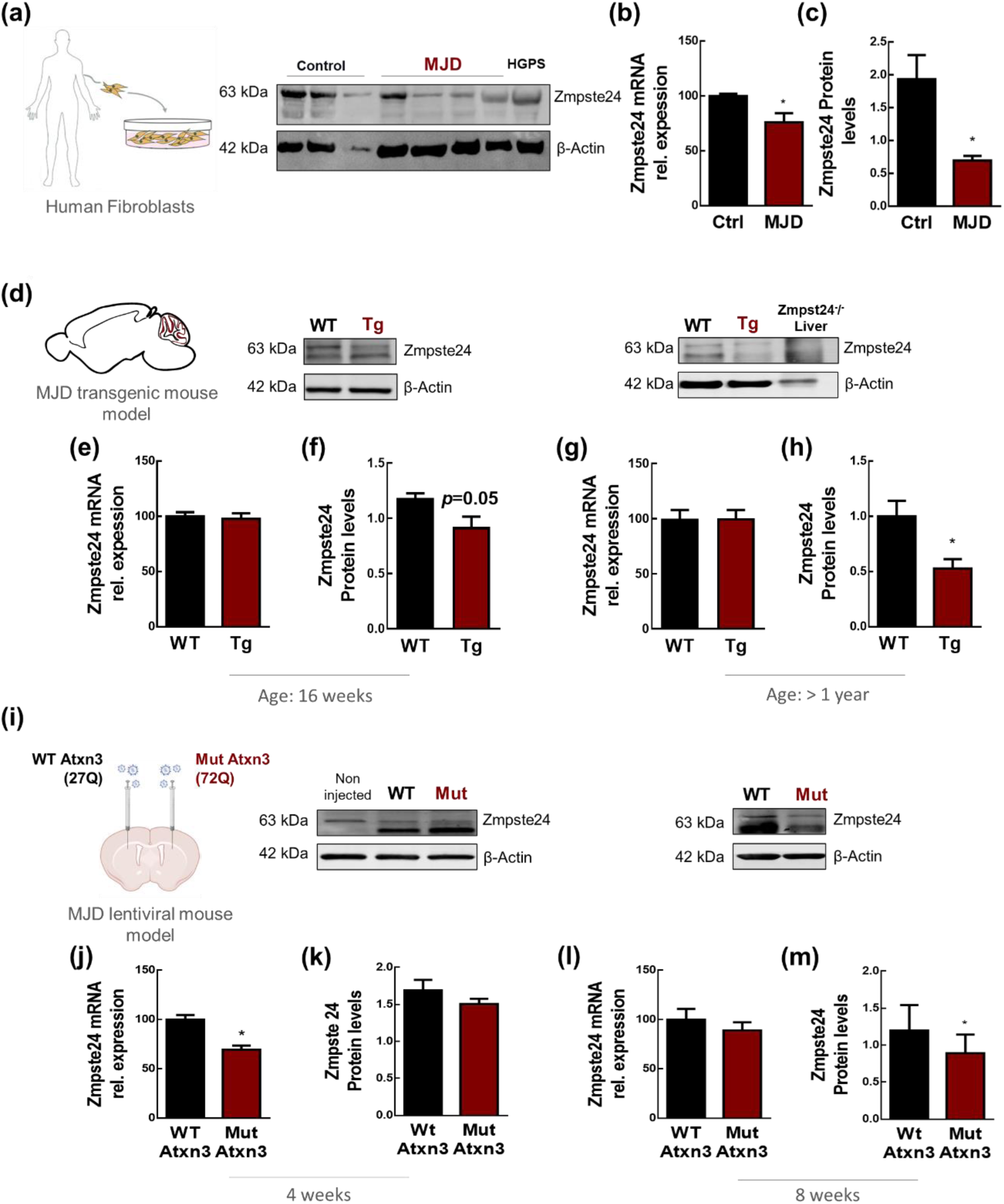
ZMPSTE24 downregulation in human and rodent MJD models. ZMPSTE24 mRNA levels evaluated by qRT-PCR, and protein levels quantified through WB in **(a)** human-derived fibroblasts, **(d)** MJD transgenic mouse model, and **(i)** MJD lentiviral-based mouse model stereotaxically injected with lentiviruses encoding for wild type ataxin-3 ((27Q) - WT Atxn3) in the left hemisphere and mutant ataxin-3 ((72Q) - Mut Atxn3) in the right hemisphere, at the following coordinates (x: + 0.6; y: +/- 1.8; z: −3.3). Tissues were collected and processed for WB and qRT-PCR analysis, after the sacrifice of MJD transgenic mice with 16 weeks and >1 year of age, and from MJD lentiviral mouse model after 4 and 8 weeks of the surgery. Significant decrease in the mRNA and protein levels of ZMPSTE24 in MJD human fibroblasts comparing with controls **(b,c)** and in the protein levels of transgenic mice with > 1 year of age **(h)** with no alterations seen at other time points in this mouse model **(e, f, g)**. Significant decrease in the mRNA levels of *Zmpste24* in MJD lentiviral model at 4-week post-injection **(j)** and protein levels at 8-week post-injection **(m)**. No alterations found in the protein levels of mice with 4 weeks (**k**) and mRNA levels in mice with 8 weeks post-injection **(1l).** All protein relative levels were normalized with β- Actin. qRT-PCR analysis was normalized with endogenous control (*GAPDH* and *Gapdh*). Statistical significance was evaluated with Unpaired t-test *p<0.05, n=3/4 (human fibroblasts); n=5 (Tg - 16 weeks); n=7 (Tg>1y); and paired Student’s t-test *p < 0.05, n = 5 (LV model). Data are expressed as mean ± SEM. MJD – Machado-Joseph disease, HGPS – Hutchinson Gilford Progeria Syndrome, WB – Western blot. WT – wild type. Last brain scheme created with BioRender.com.

The decreased levels of ZMPSTE24 have been correlated to ectopic overexpression of GMF-β in non-neuronal cells, and a possible consequence/response to oxidative stress, which appears to be associated with the activation of p53 and its downstream target gene - *P21/Waf1* (Imai et al., 2015). In line with that, we investigated the presence of those factors in a MJD lentiviral-based mouse model, but our results did not confirm these mechanisms of ZMPSTE24 downregulation in MJD, rather a significant GMF-β downregulation was found in old transgenic mice (Figure S1d) suggesting that a different mechanism might be involved. The levels of *P21/Waf1* were also found to be downregulated in old transgenic mouse model (Figure S1e). Transgenic mice with 16 weeks of age and the MJD lentiviral mouse model did not show any alterations (Figure S1b, c, g-j)

### 2.2 Nuclear lamina alterations in MJD models

The nuclear envelope is of major importance, being the central organizing element in eukaryotic cells. As already mentioned, it has crucial functions shielding the genetic material, ensuring its regulated transcription, repair and replication. Its integrity has been reported to be clearly compromised in aging and in age related disorders. Nuclear envelope’s integrity is supported by the presence of a mesh-like structure composed by intermediate filaments – lamins – which lay beneath the inner nuclear membrane (Robijns et al., 2018). To evaluate if this structure is affected in MJD, lamins A/C and B levels were assessed in MJD patient fibroblasts, as well as in the transgenic and lentiviral-based MJD mouse models, comparing with the healthy counterparts (Figure 2a, d, k). In human fibroblasts, we did not detect any differences in lamins A/C and B levels between MJD and healthy donors (Figure 2b, c). Nevertheless, a decrease of 44.7 ± 12.0% and 15.3 ± 7.2% in lamin B protein levels was observed in the cerebellum of transgenic MJD mice, in two different age-divided groups, in comparison with WT littermates (Figure 2f, i). A 9.8 ± 3.2% decrease was also seen in the lentiviral-based MJD model, 4 weeks post-injections (Figure 2m) but not in mice sacrificed 8 weeks post-injection (Figure 2p). Furthermore, we also observed a 37.7±14.9 and 24.8 ± 11.7 decrease of lamin C protein levels in the MJD transgenic mouse model, in comparison with WT littermates (Figure 2g, j) still not presenting alterations in the lentiviral mouse model (Figure 2n, q). Though not reaching significance, tendencies for decreased levels of lamin A were observed in the MJD transgenic mouse model at 16 weeks of age (Figure 2e) and in the lentiviral-based MJD model (Figure 2l, o), while older MJD transgenic mice (over 1 year of age) showed a tendency to higher levels of lamin A, in comparison to WT mice (Figure 2h).

**Figure 2.**
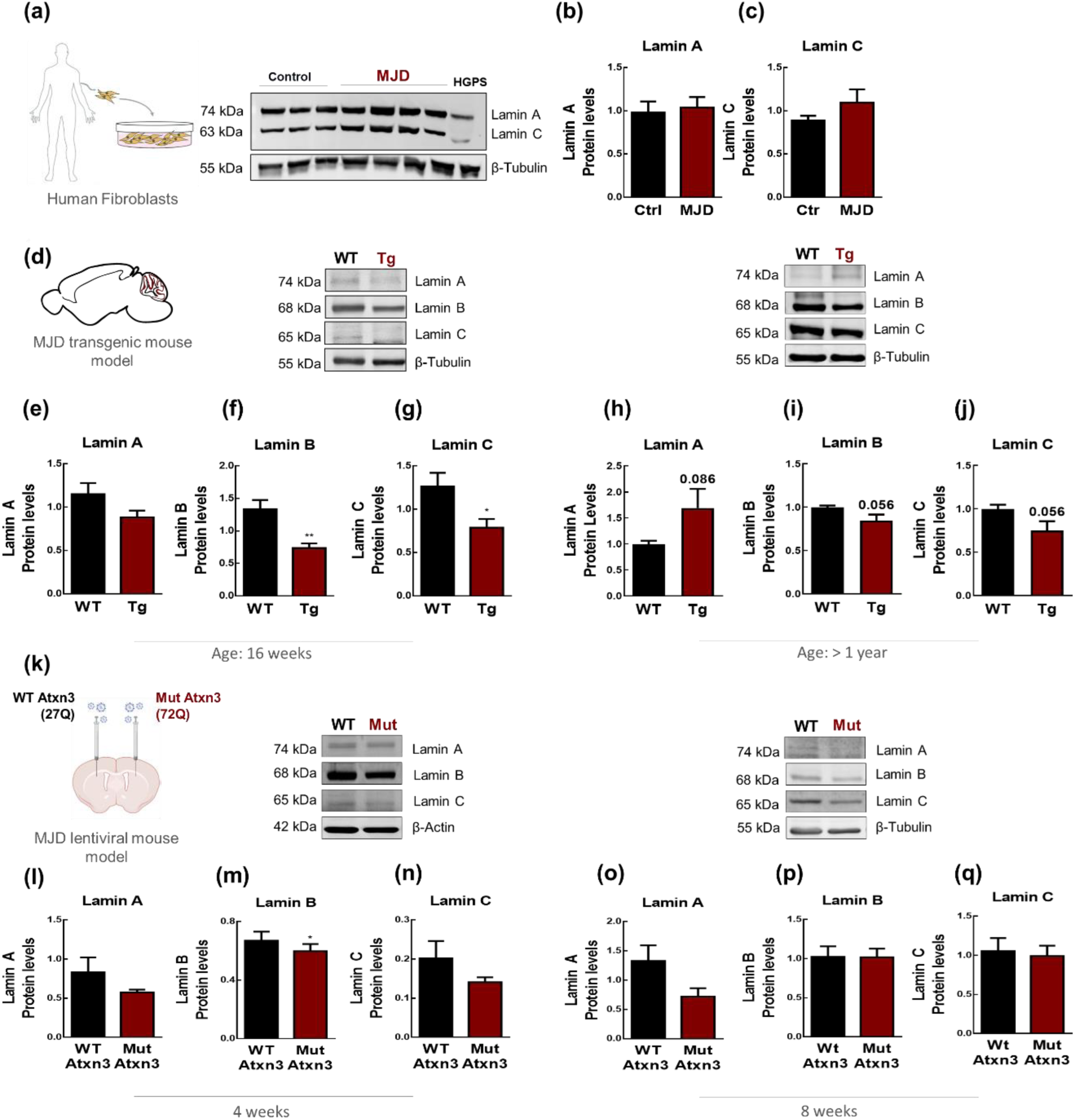
Nuclear lamina alterations in MJD models. Protein levels of lamins A/C and B quantified by WB in **(a)** human fibroblasts **(d)** MJD transgenic mouse model and **(k)** lentiviral-based mouse model of MJD, as described in Figure 1. No alterations found in human fibroblasts between MJD and healthy donors **(b, c)**. Lamin A shows a tendency to increase in the older transgenic mice group **(h)** and a tendency to decrease in transgenic mice with 16 weeks of age **(e)** and in the lentiviral-based MJD model **(l, o).** Significant decrease in lamin B protein levels in the cerebellum of MJD transgenic mice with 16 weeks of age **(f)** and in the lentiviral mouse model with 4 weeks post-injection **(m)**. A tendency to decreased levels seen in older transgenic mice **(i)** and no alteration found in 8-week lentiviral mouse model **(p).** Lamin C protein levels showing a significant decrease in the transgenic mouse model with 16 weeks of age **(g)** and a tendency to decreased levels in the remaining models **(j, n, q)** In the lentiviral-based mouse model **(k)**, a significant decrease in the lamin B protein levels is observed only at 4-week post-injection **(m)**. Statistical significance was evaluated with Unpaired t-test (*p<0.05, **p< 0.01) n=3/4 (human fibroblasts); n=5 (Tg - 16 weeks); n=7 (Tg >1year); and paired Student’s t-test (*p< 0.05) n=5 (LV-based mouse model). Data are expressed as mean ± SEM.

The loss of lamins, at the mRNA level, was also associated and suggested as a marker of cellular senescence (Freund et al., 2012). Dysfunction of the B-type lamins leads to functional defects in adult neurons, regarding chromatin formation, cell cycle activation and neuronal survival, being considered as the most important risk factor for the development of common neurodegenerative diseases and basic mechanisms of cellular senescence (Frost, 2016). With that in mind, we investigated lamins B1 and B2 mRNA levels in both MJD transgenic mice and the lentiviral-based MJD model (Figure 3a, f). We observed a significant decrease of 35.7 ± 11.2 in lamin B1 and 30.9 ± 4.9% in lamin B2 mRNA levels in the MJD transgenic mouse model at 16 weeks of age (Figure 3b, c). Nonetheless, this decrease was not sustained over time (Figure 3d, e) nor detected in the lentiviral-based mouse model (Figure 3g-j).

**Figure 3.**
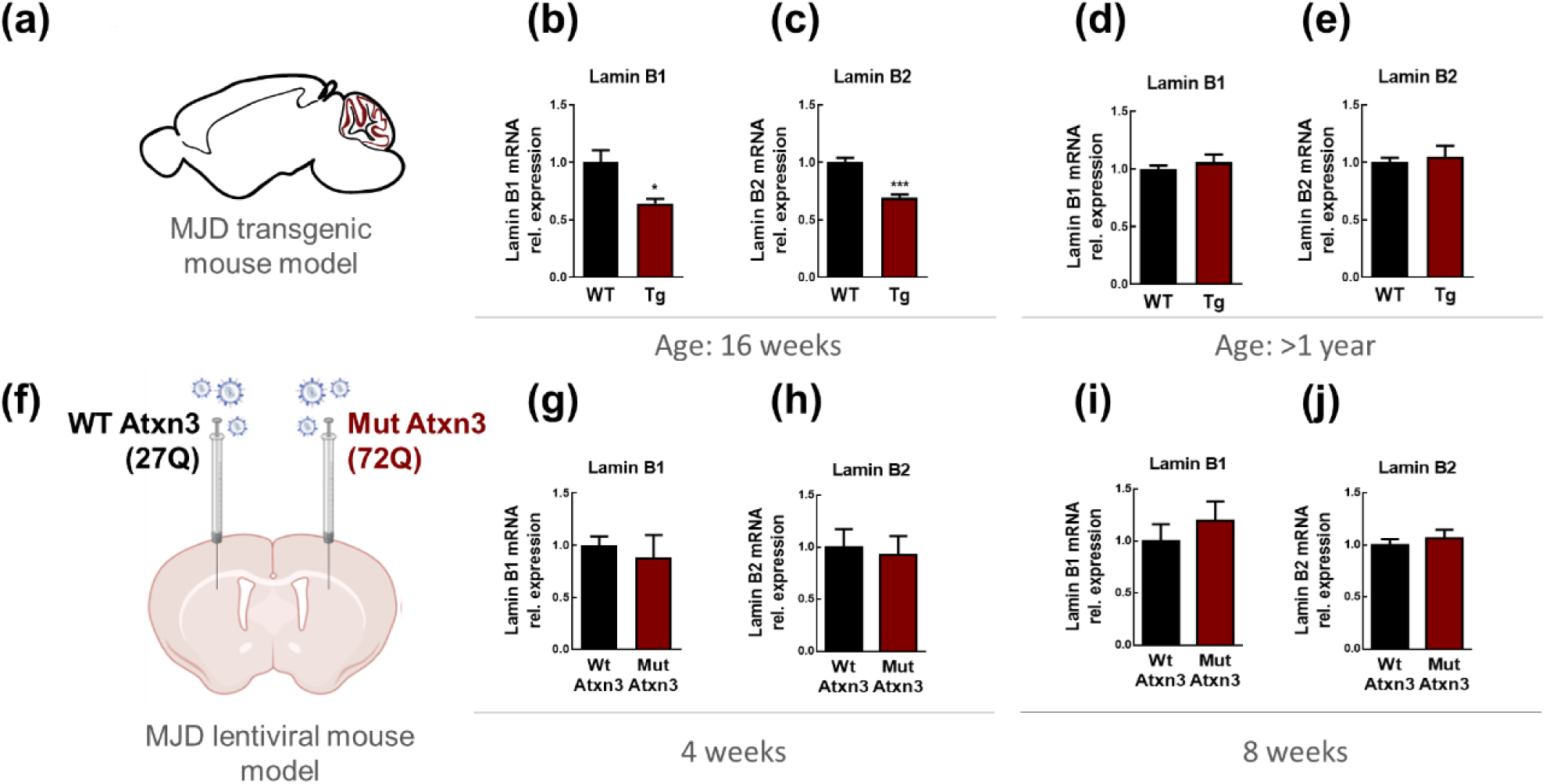
Lamin B transcriptional downregulation alteration in MJD. Lamins B1 and B2 mRNA relative expression evaluated by qRT-PCR in **(a)** MJD transgenic mouse model and **(f)** MJD lentiviral-based mouse model, submitted to the same procedures described in Figure 1. Significant decrease of lamins B1 **(b)** and B2 **(c)** mRNA levels in the cerebellum of MJD transgenic mice with 16 weeks of age and no alterations seen in older mice **(d, e)** neither in the lentiviral model **(g-j)**. qRT-PCR analysis was normalized with endogenous control (*Gapdh*). Statistical significance was evaluated with Unpaired t-test (*p<0.05, ***p< 0.001), n=5 (Tg - 16 weeks); n=7 (Tg >1year); and paired Student’s t-test (*p< 0.05) n=4 (4 weeks), n=5 (8 weeks) (LV-based mouse model). Data are expressed as mean ± SEM.

Globally, these results suggest that the levels of components of nuclear lamina are dysregulated, most of the times reduced, in MJD.

### 2.3 Loss of nuclear integrity in cellular models of MJD

Besides nuclear envelope integrity loss, another strong evidence of aging is the presence of nuclear shape abnormalities, as seen in HGPS and in normal aged cells (Carrero et al., 2016; Ragnauth et al., 2010).

To assess if mutant ataxin-3 could cause a decrease in the levels of proteins associated with nuclear integrity, and consequently induce nuclear shape abnormalities, we measured the nuclear circularity in N2a cells, after transfection with different constructs of this protein, two of them, more aggregation-prone.

The following constructs were transfected: as a control, WT Atxn3 – non-expanded ataxin-3 - LV-PGK-Atx3 27Q (Alves et al., 2008); Q72 - C-terminal fragment of mutant ataxin-3 (Carmona et al., 2017); Q69 – C-terminal fragment; N terminally-truncated mutant ataxin-3 (Torashima et al., 2008); Mut Atxn3 – full-length mutant ataxin-3 - LV-PGK-Atx3 72Q (Alves et al., 2008) (Figure 4 a).

**Figure 4.**
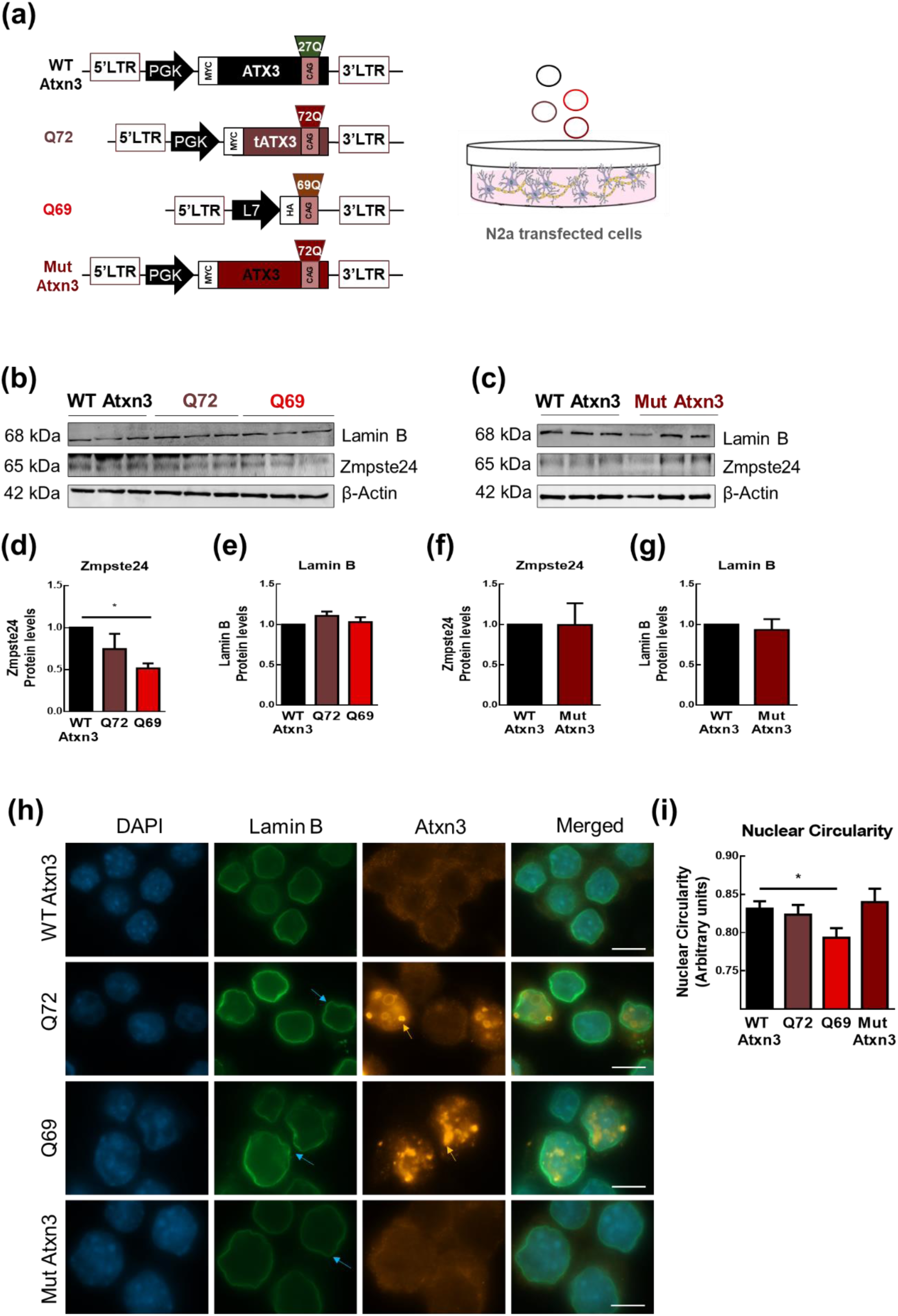
Nuclear circularity loss and dysregulation in N2a MJD cell models. **(a)** Schematic representation of the four different ataxin-3 constructs (with different aggregation properties): WT Atxn3, Q72, Q69 and Mut Atxn3 used to create N2a models of MJD by transfection. **(b, c)** Levels of ZMPSTE24 and lamin B quantified by WB **(d-g)**, displaying a significant decrease in Q69 transfected cells, comparing with WT Atxn3 **(d)**. **(h)** Immunocytochemical staining of nuclear envelope protein - lamin B of the transfected N2a MJD cell model. Orange: ataxin3; Green: lamin B; Blue: DAPI nuclear staining; Blue arrow: nuclear blebs; Orange arrow: aggregates. **(i)** Evaluation of nuclear circularity of the cells transfected with the different ataxin-3 constructs, showing a significant loss of circularity in cells transfected with the construct bearing Q69. Nuclear circularity was analysed using Fiji (Fiji Is Just ImageJ, NIH) Statistical significance was evaluated with One-way ANOVA with Dunnett’s multiple comparisons test (*p<0.05), n=3. Scale bar, 10 µm.

As shown in Figure 4h, the transfection of N2a cells with the construct that is most prone to aggregate (Q69), led to a significant decrease in ZMPSTE24 protein levels (48.44± 5.8%) (Figure 4b, d). This result might explain the more prominent modifications of nuclear shape also observed in this condition, when compared to WT Atxn3 transfected cells (Figure 4h, i) which is also reinforced by the lack of alterations induced by the other constructs (Figure 4 f, g). The aggregation of ataxin-3 apparently did not impact on lamin-B levels (Figure 4 e).

Altogether this data indicates a loss of nuclear integrity associated with the accumulation of mutant ataxin-3 aggregates.

### 2.4 Progerin overexpression aggravates neuropathology and neuroinflammation in a lentiviral-based mouse model of MJD

Our previous data suggest that MJD models present, to some extent, characteristics of an aged phenotype, displaying less expression of nuclear lamins and associated proteins, as well as nuclear shape abnormalities, which reveals nuclear frailty. Therefore, we then investigated whether the overexpression of progerin, inducing an age-accelerated process, together with MJD neuropathology, could precipitate or aggravate MJD-associated neuropathology. For this purpose, we took advantage of a lentiviral-based mouse model of MJD, generated by the localized injection of lentiviruses encoding for the mutant ataxin-3, simultaneously with progerin in the striata of mice, and mutant ataxin-3 together with EGFP in the left control hemisphere (Figure 5a) (Alves et al., 2008). This model is particularly valuable as it allows the induction of a robust neuropathology and its precise quantification, regarding a) the loss of the DARPP-32 marker, as a consequence of striatal neuronal dysfunction caused by the accumulation of mutant ataxin-3, b) the number of ataxin-3 ubiquitinated aggregates and c) cellular death quantified by the accumulation of pyknotic nuclei in specific regions of interest. Progerin overexpression was specifically induced in the striatum (Figure 5b) and induced a significant increase (44.13 ± 10.24%) in DARPP-32 volume depletion (Figure 5c).

**Figure 5.**
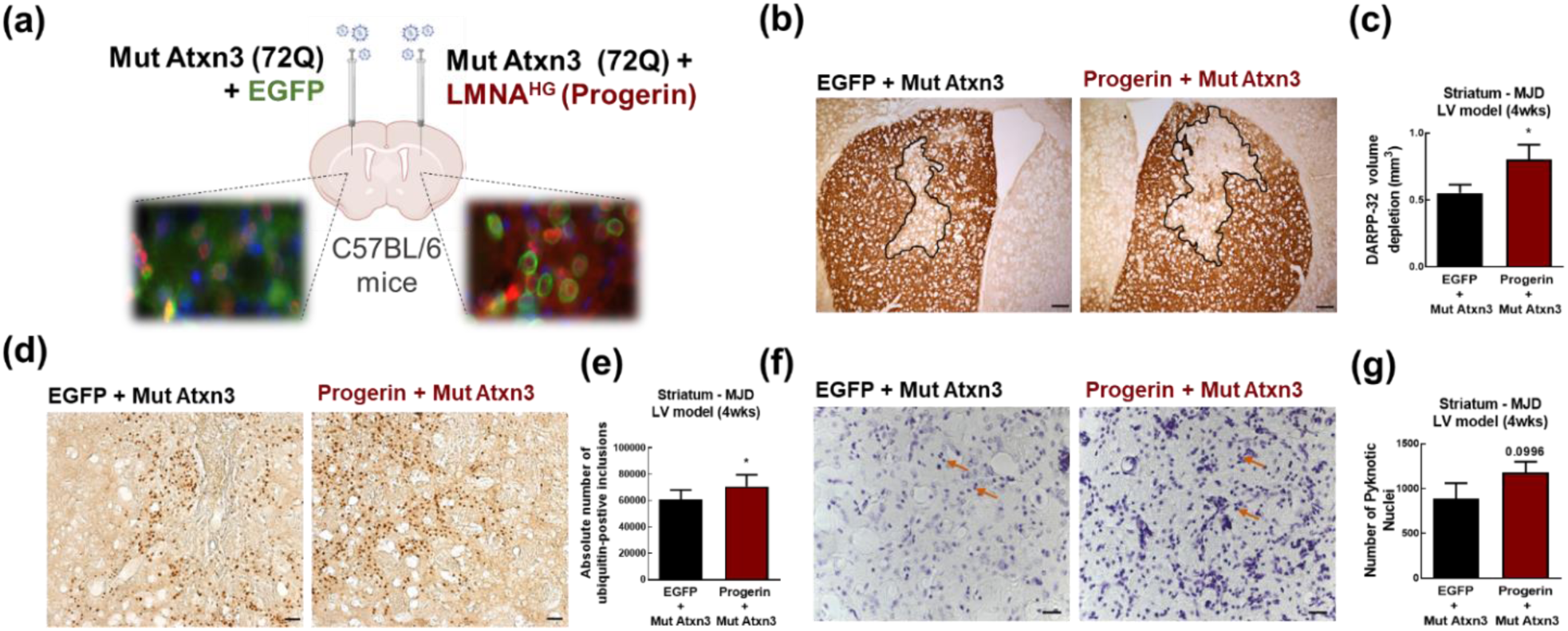
Progerin overexpression aggravates neuropathology in a lentiviral vector-based mouse model of MJD. **(a)** Schematic representation of the stereotaxic injection of lentiviruses encoding for mutant ataxin-3 (Mut Atxn3) – injected bilaterally in the striatum of six-week-old C57/BL6 mice - together with i) lentiviruses encoding for EGFP in the left control hemisphere or ii) progerin in the right hemisphere (x: + 0.6; y: +/- 1.8; z: −3.3). Mice were sacrificed at 4 and 8 weeks after surgery, and tissues processed. Detailed images showing immunofluorescence, using anti-human ataxin-3 antibody - 1H9 (Mut Atxn3 intranuclear inclusions in red), EGFP and progerin in green. **(b)** Immunohistochemical peroxidase staining using anti-DARPP32 antibody as dopaminergic loss of function marker, and depleted volume quantification **(c)**. Progerin injected hemisphere displayed a higher depletion in DARPP-32 volume when compared with the control hemisphere, 4-week post-injection. **(d)** Immunohistochemical analysis using anti-ubiquitin antibody and quantification of positive-ubiquitin inclusions, with progerin injected hemisphere displaying a significantly higher number of ubiquitinated inclusions, in 4-week post-injected mice **(e)**. **(f)** Cresyl violet staining indicating pyknotic nuclei (orange arrows), and its quantification **(g)**. Statistical significance was evaluated with paired Student’s t-test (*p < 0.05) n=4 (4 weeks), n=5 (8 weeks). Data are expressed as mean ± SEM. Scale bars represent (b) 500µm, (d) 100µm, and (f) 200µm. DARPP-32 - dopamine and cyclic AMP-regulated neuronal phosphoprotein 32.

The number of ubiquitinated inclusions (another neuropathological feature of MJD) was also measured and revealed an increase of 15.8 ± 2.5% upon progerin co-expression, 4 weeks after surgery (Figure 5d, e). At 8-weeks post-injection this effect was lost, as well as all the parameters measured (Figure S2a-h) excepting the size of aggregates, which became smaller, possibly resulting from degeneration of the cells with larger aggregates (Figure S2f).

Regarding cell death, a tendency to an increased number of pyknotic nuclei, densely stained with cresyl violet (Figure 5f), was also observed in the progerin expressing hemisphere, 4 and to a lesser extent, 8 weeks after surgery (Figure 5g).

It is also well known that during normal aging (reviewed in (López-Otín et al., 2023)) and in some neurodegenerative disorders, such as MJD, several inflammatory genes are upregulated, inducing a neuroinflammatory state in the brain, involving astrocytes and microglia, main components of neuroinflammation responses (Evert et al., 2001; Mendonça et al., 2019). Another relevant event that may be triggering neuronal dysfunction and eventually neuronal death in MJD is the accumulation of DNA strand breaks, also a hallmark of aging (López-Otín et al., 2023). This may be due to the inefficient DNA repair mechanisms, associated with the accumulation of mutant ataxin-3 in cells (Ward & La Spada, 2015). To gain insight into the role of aging on the exacerbation of the characteristic neuroinflammation observed in different MJD mouse models as well as the increased DNA damage observed in normal aging, we focused on the previous lentiviral-based mouse model where progerin and mutant ataxin-3 were co-expressed in striata of mice. Neuroinflammation was evaluated by means of immunofluorescence, using antibodies against GFAP and AIF1/Iba1(Gonçalves et al., 2013).

In the age-accelerated hemisphere, we observed a significant increase of 31.0±9.5 % in GFAP immunoreactivity, suggestive of astrocytic activation, at 4-weeks post-injection (Figure 6d, e), significance that was lost at 8-weeks post injection (Figure S3d, e). Other parameters of neuroinflammation, such as Iba1 immunoreactivity and microglial recruitment, displayed a tendency to increase in the progerin injected hemisphere at 4-week post-injection (Figure 6a-c), and no alterations were found at 8 weeks post-injection (Figure S3a-c).

**Figure 6.**
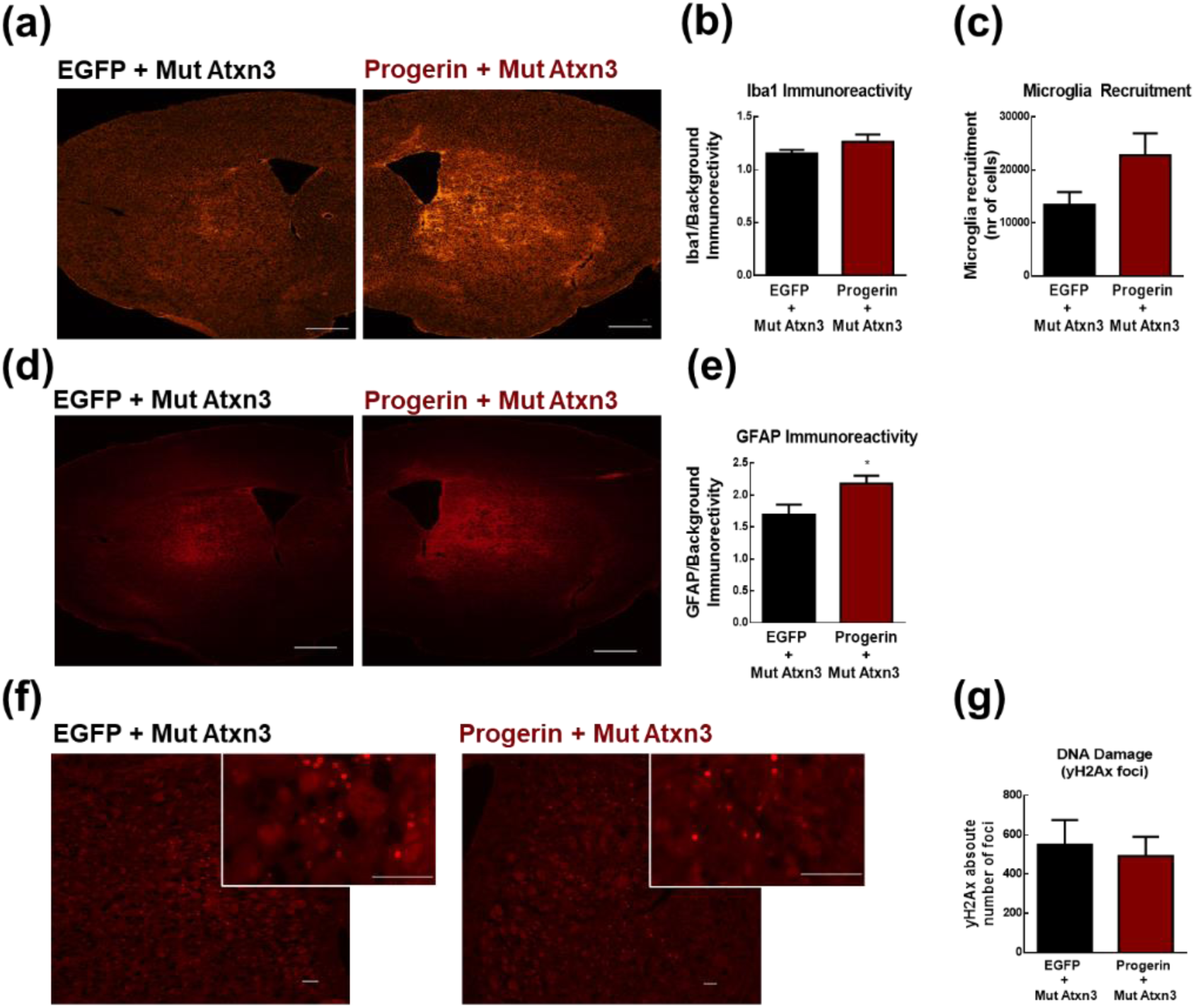
Progerin overexpression and its role in neuroinflammation and in DNA damage (4 weeks post-injection). Mice were submitted to the same procedure described in Figure 5. **(a)** Fluorescent immunohistochemical analysis of Iba-1, **(b)** relative fluorescence immunoreactivity measurement among hemispheres and **(c)** microglial recruitment (displaying a tendency to increase in the progerin-injected hemisphere at 4-week post-injection. **(d)** Fluorescent immunohistochemical analysis of GFAP in the striatum, showing a significant increase in the progerin-injected hemisphere **(e)**. **(f)** Fluorescent immunohistochemistry for the detection and quantification of DNA damage using the phosphorylation of γH2AX as a marker/antibody, presenting no alterations among hemispheres **(g)**. Microglial recruitment and γH2AX foci measured using Fiji (Fiji Is Just ImageJ, NIH). Statistical significance was evaluated with paired Student’s t-test (*p < 0.05) n=4. Data are expressed as mean ± SEM. Scale bars represent **(a,d)** 500µm, (f) 100µm. Iba-1 – ionized calcium binding adaptor molecule 1; GFAP-glial fibrillary acidic protein.

To investigate DNA damage, progerin and mutant ataxin-3 overexpressing brain sections were stained with an antibody for phosphorylated γH2AX, as previously described (Redon et al., 2009). We observed that ataxin-3 induced the formation of γH2AX foci, in accordance with earlier reports (Ward & La Spada, 2015). Interestingly, despite not being able to observe differences between age-accelerated and control hemispheres (Figure 6f, g and Figure S3f, g)), a tendency to increased protein levels of γH2AX was found in cerebellar nuclear fractions of the transgenic mouse model (S4).

Taken together, these results suggest that aging, induced by progerin, aggravates MJD pathogenesis, intensifies MJD-associated neuroinflammation and may exert some impact in DNA damage as well.

## 3 DISCUSSION

Polyglutamine diseases, including MJD, are typical late age-of-onset diseases, with symptoms appearing in midlife, and characterized by nuclear inclusions of the abnormally expanded polyglutamine proteins. Despite the growing evidence of the important role of aging in the pathogenesis of such diseases, the molecular mechanisms responsible for this late onset, are still not well understood, despite being considered the principal risk factor for the onset of several neurodegenerative disorders associated with aggregate depositions, such as Alzheimer’s, Huntington’s and Parkinson’s disease (Knowles et al., 2014; Mattson & Magnus, 2006). Indeed, it was already shown that fragility of striatal neurons to mutant huntingtin increases with age (Diguet et al., 2009).

Having the main hallmark of MJD in mind - the accumulation of intranuclear inclusions - we hypothesized that the role of aging in this pathology would also be related with nuclear integrity. We have shown the presence of age-related markers associated with nuclear integrity in cellular and animal models of MJD and, inducing an age-accelerated context into MJD pathogenesis (through the overexpression of progerin), we were able to induce an aggravation in MJD neuropathology and associated neuroinflammation, suggesting a significant role of aging in MJD pathogenesis.

Given the association of this aging-associated nuclear markers, not only with normal aging, but also with neurodegenerative disorders, we focused our research on the nuclear envelope components, particularly the lamina content and their associated processing protein, ZMPSTE24 (Carrero et al., 2016; Chang et al., 2011; Frost, 2016; Ragnauth et al., 2010). Therefore, we investigated the presence of these markers in different cellular (patient fibroblasts, N2a cells) and animal models (transgenic and lentiviral model) of MJD.

We found a significant reduction in both mRNA and protein levels of ZMPSTE24, in different MJD models, namely human fibroblasts, transgenic and lentiviral-based mouse models, suggesting a compromised expression of ZMPSTE24 in MJD. Moreover, we found a tendency to increased levels of lamin A in old transgenic mice, a drastic decrease in lamin B in two different models of MJD, and a decrease in lamin C levels, comparing with their respective healthy counterparts. These observations suggest that, in fact, nuclear integrity may be compromised in MJD, given the abnormal levels of these essential elements. Confirming this reasoning, we found an increased number of nuclei presenting abnormal shape, another “hallmark” of aging shared with progeroid syndromes (Carrero et al., 2016) upon mutant ataxin-3 (Q69) overexpression. Additionally, a significant decrease in ZMPSTE24 protein levels was observed, which may probably be one of the causes for the abnormal nuclear shape.

As future perspectives, we are willing to explore which mechanisms are underlying the dysregulation of the levels of lamins and ZMPSTE24. The decreased levels of lamin B1 in Alzheimer’s disease have already been linked to over-stabilization of filamentous actin, causing clustering of the linker of nucleoskeleton and cytoskeleton (LINC) complex along the nuclear envelope, leading to a reduction in B-type lamin protein levels (Frost, 2016). However, it remains unclear whether the cellular neurodegenerative context in MJD would be similar to the one created by the accumulation of tau. Nevertheless, it was already shown that ataxin-3 interacts with α-tubulin and binds microtubules in human, mouse and *C. elegans,* with a peculiar distribution during cell cycle, important to a correct cell division through an intact microtubule network, and its absence leads to cells’ ubiquitylated foci accumulation, cytoskeletal (including actin cytoskeleton networks) and adhesion (through downregulation of α5 integrin signalling) defects and increased cell death (Neves-Carvalho et al., 2015).

The decreased levels of ZMPSTE24 were also found to be correlated to ectopic overexpression of GMF-β in non-neuronal cells, possibly as consequence/response to oxidative stress, which appears to be associated with the activation of p53/p21 (Imai et al., 2015). More importantly, GMF-β has been found to be upregulated in several neuroinflammation and neurodegeneration conditions (Fan et al., 2018). Our results, however, did not confirm these mechanisms of ZMPSTE24 downregulation in MJD (Figure S1). Rather, we did find a significant downregulation of GMF-β in old transgenic mice (Figure S1 d). Despite its high expression being correlated with the inflammatory status of several neurodegenerative disorders, it has been shown that the absence of GMF led to impaired motor activity and motor learning, in particular beam walking and eyeblink conditioned response, being correlated with the histologic finding of neuronal loss in the inferior olive. These findings, however, are mainly relating the impact of GMF absence at, as early as, the developmental stage (Lim et al., 2004). We can speculate that its downregulation might also have an impact in the adulthood, however more evidence is needed to support this hypothesis. Furthermore, p21/waf1 was also found to be downregulated in old transgenic mice (Figure S1 e), which goes in accordance with previous studies (Imai et al., 2015) and more recently a study showing Cdk1α (*P21/Waf1*) downregulation in pre-symptomatic Huntington’s transgenic mouse model (9 months), more specifically in cerebellar Gad2^+^ neurons, despite the vast majority pointing out an upregulation associated with aging, becoming also an interesting result to be studied and pursued in our models of this specific polyglutamine disorder (Bauer et al., 2023).

Polyglutamine-expanded ataxin-3 was shown to retain enhanced interaction and deubiquitinating catalytic activity of p53, causing a more severe p53-dependent neurodegeneration in zebrafish brains and in substantia nigra and striatum of a transgenic MJD mouse model (Liu et al., 2016). Other pathways should be explored to clarify this mechanism. Aggregates found in polyglutamine diseases have been implicated in the sequestration of several transcription factors in aggresomes, such as TATA-binding protein (TBP), TBP-associated factor (TAF (II) 130), Sp1, adenosine 3′,5′- monophosphate (cAMP) responsive element-binding protein (CREB) and CREB-binding protein (CBP) (Yamada et al., 2000). Consequently, there is a reduction in the levels of several proteins, and further neurodegeneration. The impact of such transcription factors in the reduction of lamins and ZMPSTE24 expression levels should be addressed. Another hypothesis relies in the fact that autophagy is also compromised in Machado-Joseph disease, being itself an impacting factor in the levels of lamins and associated proteins, consequently affecting the integrity of the nuclear envelope (Aveleira et al., 2020; Onofre et al., 2016).

Given the evidence of implication of aging in MJD, we used a novel approach to induce an aged context to MJD, by overexpressing progerin in the striatum of a lentiviral-based mouse model of MJD. Progerin is a mutant form of the lamin A protein, resulting from a point mutation, that is the cause of the HGPS, a syndrome characterized by the premature and accelerated aging of its carriers (Eriksson et al., 2003). The purpose was to investigate if some specific markers of aging, such as the accumulation of progerin and loss of nuclear integrity could also aggravate MJD (Carrero et al., 2016). Remarkably, progerin aggravated MJD-associated neuropathological markers, given by the increased loss of DARPP-32 expression, increased number of ubiquitinated aggregates, together with a significant increase in reactive astrogliosis. However, this aggravation was not accompanied by increased accumulation of DNA damage evaluated by immunostaining of γH2AX foci. This suggests that the loss of integrity of the nuclear envelope in this specific model, aggravates the disease by further promoting more accumulation of ubiquitinated aggregated species, which induces a more immunoreactive state, independent of DNA damage. Taking all the above-mentioned evidence into account, we may infer that aging may play an important role in the aggravation of MJD pathogenesis and associated neuroinflammation.

In summary, this work dissected the impact of aging mechanisms on MJD course. Having in mind that by accelerating aging, MJD-associated neuropathology can be aggravated, a possible strategy to delay neurodegeneration can rely on delaying aging itself. This statement can be supported by our previous studies, where beneficial effects were observed using anti-aging strategies, such as caloric restriction, the usage of resveratrol or the overexpression of neuropeptide Y in MJD models (Cunha-Santos et al., 2016; Duarte-Neves et al., 2016). Furthermore, the decreased levels of ZMPSTE24 or lamins B and C observed in both cellular and animal models of MJD can be potential candidates, not only as potential therapeutic targets, but also as biomarkers of the disease.

## 4 EXPERIMENTAL PROCEDURES

### 4.1 Human fibroblast culture

MJD and control fibroblasts were generated from dermal biopsies following informed consent, under protocols approved by the Ethics Committee of the Medical Faculty of the University of Coimbra, as described in (Onofre et al., 2016). Unaffected control individuals: female, 44 years; male, 47 years; female, 52 years. Machado-Joseph disease patients: male, 22 years, 80 CAG; female, 25 years, 77 CAG; female, 31 years, 79 CAG; male 69 years, 70 CAG.

Cells were maintained in culture using high glucose Dulbecco’s modified Eagle’s medium (DMEM, Sigma-Aldrich, St. Louis, USA) supplemented with 10% foetal bovine serum (FBS, Life Technologies), 1% nonessential amino acids, 2 mM L-glutamine (Life Technologies), 1% penicillin/streptomycin (Life Technologies), 110 mg/L sodium pyruvate (Sigma-Aldrich) and 44 mM sodium bicarbonate (Sigma-Aldrich) and kept at 37°C in a 5% CO_2_ atmosphere incubator.

### 4.2 Culture and transfection of human and murine cellular lines

Human embryonic kidney 293, stably expressing the SV40 large T antigen (HEK293T) and mouse neuroblastoma cell line (N2a) obtained from the American Type Culture Collection cell biology bank (CCL-131), were maintained in standard DMEM medium, supplemented with 10% FBS and 1% penicillin/streptomycin at 37°C under a humidified atmosphere containing 5% CO_2_.

For the transfection of N2a cell line, 1×10^5^ cells were plated on a 12-well cell culture multiwell plate containing poly-D lysine-coated coverslips (Sigma-Aldrich). After 24 hours, medium was reduced to half and cells were transfected with a mixture of DNA/polyethyleneimine (PEI) complexes (MW40000, PolySciences). Complexes were formed by combining 500 ng of plasmid DNA in 10 μL of serum free DMEM and 3 μL of PEI (1 mg/ml) per well. DNA/PEI were then diluted in complete DMEM and added to cells. The transfected vectors were: wild-type ataxin-3 (WT Atxn3 - LV-PGK-Atx3 27Q) and mutant ataxin-3 (Mut Atxn3 - LV-PGK-Atx3 72Q) (Alves et al., 2008); N-terminal-truncated human ataxin-3 with 69 repeats and a N-terminal hemagglutinin (HA) (Q69 – LV-L7-TmutATXN3 69Q) (Torashima et al., 2008) and C-terminal fragment of mut Atxn3 with 72 glutamines (Q72 - LV-PGK-TmutATXN3 72Q) (Carmona et al., 2017).

Cells were fixed 48 hours post-transfection with 4% paraformaldehyde (PFA, Sigma-Aldrich) for 20 minutes at RT, washed with PBS and kept at 4°C until further use.

### 4.3 Lentiviral production, purification, and titer assessment

Lentiviral vectors encoding for EGFP (LV-PGK-EGFP), WT Atxn3 (LV-PGK-Atx3 27Q), Mut Atxn3 (LV-PGK-Atx3 72Q) (Alves et al., 2008) and Progerin (LMNA^HG^ - pCDHblast MCSNard OST-LMNAd50 – kindly provided by Professor Tom Misteli (Addgene plasmid #22662) (Pegoraro et al., 2009) were produced in HEK293T cells with a four-plasmid system, as previously described (de Almeida et al., 2002). Lentiviral particles were resuspended in sterile 0,5% bovine serum albumin (BSA, Millipore) in phosphate buffered solution (PBS). The viral particle content was assessed by measuring HIV-1 p24 antigen levels (Retro Tek, Gentaur). Viral concentrated stocks were kept under - 80°C until further use.

### 4.4 Animal experimentation

#### 4.4.1 Animal maintenance

C57Bl/6-background transgenic MJD mice and WT control littermates with either 16 weeks or 1 to 2 years of age, were obtained by backcrossing heterozygous males with C57Bl/6 females (obtained from Charles River, Barcelona, Spain). Transgenic mice were acquired from Hirokazu Hirai, Kanazawa University, Japan (Torashima et al., 2008) and a colony of these transgenic mice was established in the licensed animal facility (International Animal Welfare Assurance number 520.000.000.2006) at the Center for Neuroscience and Cell Biology of the University of Coimbra (CNC-UC). Transgenic mice express specifically in cerebellar Purkinje cells a N-terminal truncated human ataxin-3 with 69 repeats and a N-terminal hemagglutinin (HA) epitope, driven by a L7 promotor. Genotyping was performed by means of PCR, using DNA from ear tissue.

Five-week-old C57Bl/6 mice, obtained from Charles River (Barcelona, Spain) were also used and submitted to striatal stereotaxic surgery.

Mice were housed under conventional 12-hour light/dark cycle in a temperature-controlled room with water and food *ad libitum*. The experiments were carried out in accordance with the European Union Community directive (2010/63/EU) for the care and use of laboratory animals. Researchers received adequate training (FELASA certified course) and certification to perform the experiments from the Portuguese authorities (Direcção Geral de Veterinária) (ORBEA_289_2021/10122021).

#### 4.4.2 Mouse surgery

Six-week-old C57BL/6J mice were anesthetized by intraperitoneal administration of avertin (14 μL/g, 250 mg/Kg body weight). Mice were stereotaxically injected into the striatum with the following coordinates, calculated from *bregma*: anteroposterior: +0.6 mm; lateral: ±1.8 mm; ventral: −3.3 mm; tooth bar: 0. A single injection of concentrated lentiviral vectors containing 400 ng of p24 antigen of each vector encoding for the specific transgenes, was given on the corresponding hemispheres, at an infusion rate of 0.25 μL/min using a 10 μL Hamilton syringe.

Concentrated viral stocks were thawed on ice, and diluted viral suspensions were prepared using sterile 0,5% BSA in PBS.

In a group of animals, lentiviral vectors encoding for either the wild-type ataxin-3 (Atx3 27Q) in the left hemisphere or the mutant ataxin-3 (Atx3 72Q) in the right hemisphere were stereotaxically injected in the striatum. After surgery, mice were maintained in their home cages and were sacrificed 4 or 8 weeks later for western blotting and qPCR analysis.

Another group of animals was injected with lentiviruses encoding for the mutant ataxin-3 (72Q) and EGFP in the left hemisphere, or the mutant ataxin-3 (72Q) and Progerin (LMNA^HG^) in the right hemisphere. After surgery, mice were maintained in their home cages and were sacrificed 4 or 8 weeks later for immunohistochemical analysis.

### 4.3 Isolation of total RNA from mouse tissue and cells

Animals were sacrificed with a lethal dose of Xylazine/Ketamine and cervical dislocation. Cerebellum or striatum of mice were dissected and stored at −80°C until further processing.

For transgenic mouse tissue, RNA was isolated with NucleoSpin RNA II isolation kit (Machery Nagel) according to the manufacturer’s instructions, after a homogenization step of the tissue with TRIzol™ Reagent (Invitrogen) and d-chloroform.

For the lentiviral mouse model, the total RNA content from tissues was isolated using the miRCURY™ RNA Isolation Kit (Exiqon), following the manufacturer’s instructions.

RNA from cells was isolated using NucleoSpin RNA II isolation kit (Machery Nagel), in accordance with the manufacturer’s instructions, protocol that was preceded by homogenization and lysis steps using kit’s Lysis Buffer, supplemented with with β-mercaptoethanol.

The amount of RNA was quantified by optical density (OD) using a Nanodrop 2000 Spectrophotometer (Thermo Fisher Scientific). RNA was stored at −80°C until further use.

### 4.4 cDNA synthesis and quantitative Real-Time Polymerase Chain Reaction (qRT-PCR)

cDNA was synthesized by conversion of 1 µg of the DNAse-I treated RNA with iScript Selected cDNA Synthesis kit (Bio-Rad), according to the manufacturer’s instructions, and stored at −20°C.

qRT-PCR was performed in the StepOne Plus Real-Time PCR System (Applied Biosystems) with SSO Advanced SYBR Green Supermix (Bio-Rad). PCR was carried out in 10 μL reaction volume using 500 nM of each primer. Primers for mouse *Zmpste24*, mouse *Lmnb1* and *Lmnb2*, human and mouse *GAPDH* were designed using Primer Blast Software (Ye et al 2012) (oligonucleotide sequences in Table 1). Primers for human *ZMPSTE24* Hs_ZMPSTE24_1_SG QuantiTect) were predesigned and validated by QIAGEN and used diluted to a final work solution of 1x (QuantiTect Primers, QIAGEN). Appropriate negative controls were also prepared. All reactions were performed in duplicate with the following running method specifications: 95°C for 30 seconds, followed by 45 cycles at 95°C for 5 seconds, and 59°C to 62°C for 15 seconds and 0.5°C increment for starting at 65°C at each 5 sec/step up to 95°C. The amplification rate for each target was evaluated from the cycle threshold (Ct) determined automatically by the Software with cDNA dilutions, relative to *GADPH* levels. The mRNA fold increase or fold decrease with respect to control samples was determined by the Pfäffl method.

**Table 1.**
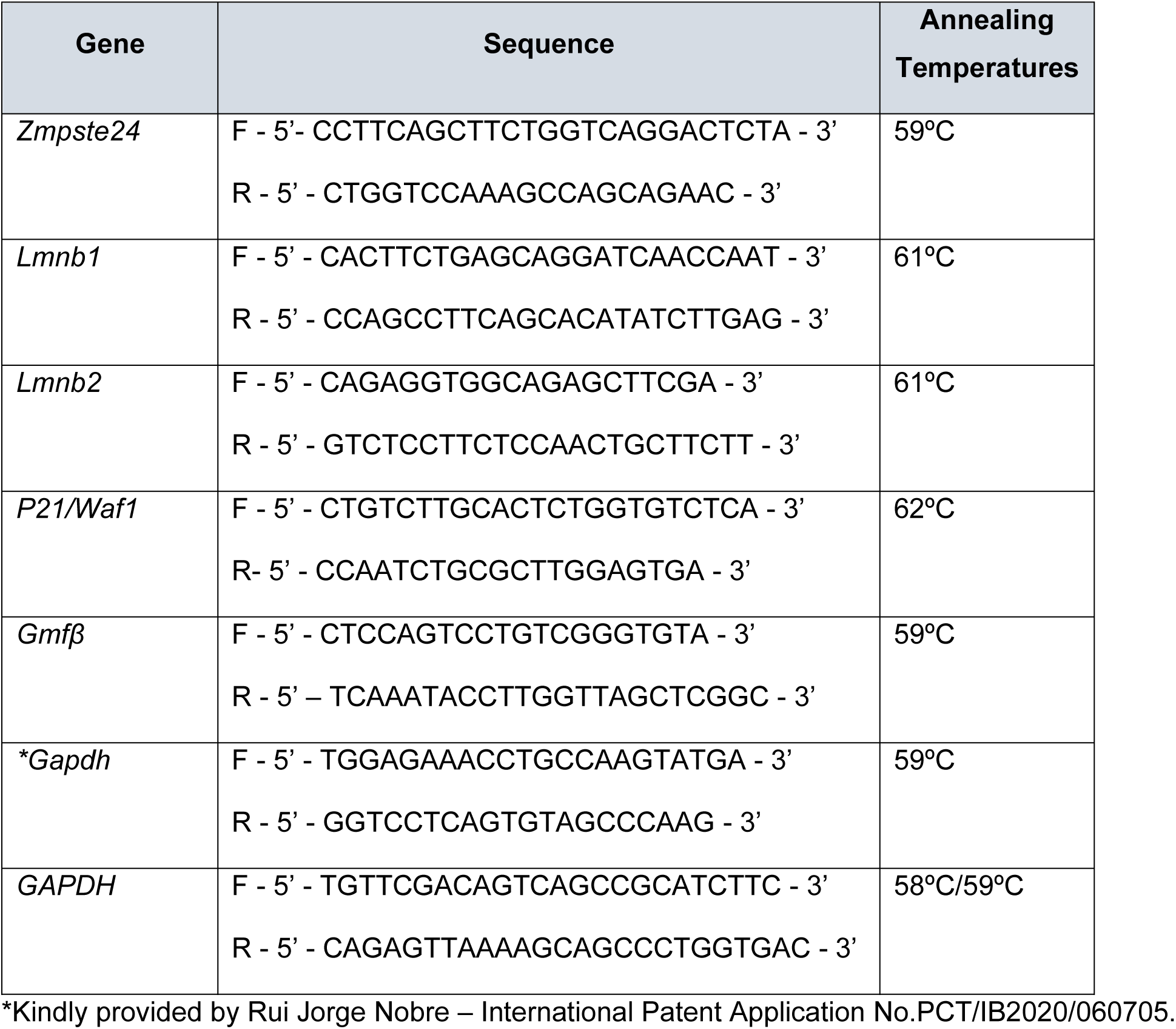
Primer Sequences

### 4.5 Protein extraction and Western Blot

Cerebellar or striatal tissues were dissected and further lysed with radioimmunoprecipitation assay-buffer solution (RIPA buffer: 50 mM Tris-base (Fisher Scientific GmbH); 150 mM NaCl (Fisher Chemical); 5 mM EGTA (ethylene glycol tetraacetic acid, Sigma-Aldrich); 1% Triton X100 (Sigma-Aldrich); 0.5% sodium deoxycholate (Sigma-Aldrich); 0.1% SDS (sodium dodecyl sulphate, Acros Organics)) containing protease inhibitors (Roche Diagnostics GmbH) and 0.2 mM PMSF (phenylmethylsulphonyl fluoride, Sigma-Aldrich), 1 mM DTT (dithiothreitol, Sigma-Aldrich, St. Louis, USA), 1 mM Sodium Orthovanadate (Sigma-Aldrich) 5 mM Sodium Fluoride (Sigma-Aldrich) and 2 mg/mL Pepstatin A (Sigma-Aldrich).

Protein concentration was determined using the Bradford protein assay (BioRad), according to the manufacturer’s instructions.

For γH2AX protein level analysis, a protein fractioning from tissues protocol was applied, using Subcellular Protein Fractionation Kit for Tissues (Thermo Scientific) according to manufacturer’s specifications.

Samples were then denatured with 6x sample buffer (9.3% DTT; 10% SDS; 30% glycerol in 0.5 M Tris-HCl/0.4% SDS - pH 6,8 and bromophenol blue (0.012%)) and incubated at 95°C for 5 minutes.

Sixty micrograms of protein extract were resolved in SDS-polyacrylamide gel (4% stacking, 10% running). The proteins were then transferred to PVDF membranes (Millipore) according to standard protocols, for 2h30 at 1 A and 4°C. After blocked in 5% nonfat milk powder in 0.05% Tween 20 – Tris-buffered saline (TBS-T) for 1 hour at RT, membranes were then incubated overnight (ON) at 4°C, with the following primary antibodies diluted in blocking solution (3% nonfat milk): goat polyclonal anti-lamin A/C (1:500; sc-6215 Santa Cruz Biotechnology); mouse monoclonal anti-lamin A/C (1:500; MANLAC1 (4A7), kindly provided by Dr Glenn Morris, Center for Inherited Neuromuscular Diseases) (Manilal et al., 2004); goat polyclonal anti-lamin B (1:750; sc-6216 Santa Cruz Biotechnology); mouse monoclonal anti-ZMSPTE24 (1:500; kindly provided by Professor Carlos Lopez-Otín) (Pendás et al., 2002); Mouse monoclonal anti-γH2AX (Ser139) clone JBW301 (1:1000, 05-636-I, Millipore); Mouse monoclonal anti-β-Actin (1:5000; A5316, clone AC74, Sigma-Aldrich); mouse monoclonal anti-β-Tubulin antibody (1:10,000; SAP.4G5 Sigma-Aldrich); rabbit monoclonal anti-TBP (1:2000, Cell signalling diluted in 5% BSA). Extended methods available in supporting information.

### 4.6 Histological staining and immunohistochemical analysis

#### Tissue preparation

Mice were sacrificed with an overdose of avertin (625 mg/kg body weight, intraperitoneally). Perfusion with PBS and fixation with 4% paraformaldehyde (PFA) were performed transcardially and brains were collected and post-fixed in 4% PFA for 24 hours. The brains were then cryoprotected/dehydrated by immersion in 25% sucrose/PBS for 36 to 48 hours. Dried brains were frozen at −80°C and 25 μm coronal sections from stereotaxically injected mice were sliced using a cryostat at −21°C. Slices were then collected in anatomical series and stored as free-floating sections in PBS supplemented with 0.05% (m/v) sodium azide.

#### Cresyl violet staining

Dried pre-mounted sections in gelatine-coated slides were dehydrated, cleared with xylene, rehydrated, and stained with the basic dye cresyl-violet solution for 3 minutes. Staining was then differentiated in 70% ethanol and sections were further dehydrated twice by passing through 95%, 100% ethanol solutions and ultimately by xylene solution, before mounting. Staining was visualized, and images captured with Zeiss Axioskop 2 imaging microscope (Carl Zeiss MicroImaging microscope, Carl Zeiss MicroImaging) equipped with AxioCam HR colour digital cameras (Carl Zeiss Microimaging) and Plan-Neofluar 5x/0.15 Ph 1 (440321), Plan-Neofluar 20x/0.50 Ph 2 (1004-989), Plan-Neofluar 40x/0.75 Ph 2 (440351) and Plan-Neofluar 63x/1.25 Oil (440460-0000) objectives using the AxioVision 4.7 software package (Carl Zeiss Microimaging).

#### Bright-field Immunohistochemistry

Free-floating brain coronal sections were treated with a 0.1% phenylhidrazyne/PBS solution, for blockage of endogenous peroxidases. After permeabilization in PBS/0.1% Triton X-100 with 10% normal goat serum (NGS), sections were incubated ON at 4 °C in blocking solution with the primary antibodies: rabbit polyclonal anti-ubiquitin antibody (1:300; ADI-SPA-200-F, Enzo Life Sciences), rabbit anti-dopamine and cyclic AMP-regulated neuronal phosphoprotein 32 (DARPP-32) antibody (1:1000; AB10518, Millipore). Sections were then washed and incubated with the biotinylated secondary goat anti-rabbit antibody (1:200 BA-1000, Vector Laboratories) 2 hours at RT. Further procedures available in supporting information

### 4.7 Immunofluorescent analysis

#### Fluorescent immunocytochemistry

Cells were fixed in PFA 4% for 15 minutes and permeabilized for 10 minutes at RT with PBS containing 0.1% Triton X-100. To block unspecific binding, cells were then incubated with PBS/10% BSA for 1 hour before incubation with primary antibodies: rabbit anti-HA (1:200, AB9110, Abcam); mouse anti-Myc-tag, clone 4A6 (1:200, 05-724, Millipore) and goat anti-lamin B (1:100, sc-6216 Santa Cruz Biotechnology) ON at 4°C, in PBS containing 3% BSA. Extended methods available in supporting information.

#### Fluorescence Immunohistochemistry

Free-floating sections were incubated in PBS/0,1% Triton X-100 containing 10% NGS and then incubated ON at 4°C in blocking solution with primary antibodies: mouse monoclonal anti-glial fibrillary acidic protein (GFAP) antibody, clone (GA5) (1:500; IF03L, Millipore); rabbit anti-ionized calcium binding adaptor molecule 1 (Iba-1) antibody (1:1000; 019-19741, Wako Chemicals, USA); mouse monoclonal anti-ataxin-3 antibody, clone 1H9 (1:5000; MAB5360, Millipore). Extended methods available in supporting information.

#### Fluorescence Immunohistochemistry - yH2AX

Sections mounted in Superfrost Plus™ Adhesion Microscope Slides (Menzel-Glaser, Thermo Scientific), underwent antigen retrieval for 30 minutes at 98°C in citrate buffer (10 mM citrate, Sigma-Aldrich) 0.05% Tween 20 (pH 6.0, Fisher Scientific GmbH) and then allowed to cool-down to RT. After washing the sections in PBS, blocking was performed for 1 hour at RT with PBS/0.3% Triton X-100 with 10% NGS. Sections were then incubated 36 hours at 4°C with the primary antibody – anti-γH2AX (Ser139) clone JBW301 (1:500, 05-636-I, Millipore) diluted in blocking buffer. Extended procedures available in supporting information.

### 4.8 Imaging Analysis

Full imaging analysis are described in the supporting information.

### 4.9 Statistical Analysis

Statistical analysis was performed with paired or unpaired Student’s t-test and One-way ANOVA with Dunnett’s multiple comparisons test, as mentioned in the respective panel of results. Results are expressed as mean ± standard error of the mean (SEM). Significant thresholds were set at p<0.05 and p<0.01, as defined in the text. Calculations were performed using GraphPad Prism version 6.00 and 8.0.1 for Windows.

## AUTHOR CONTRIBUTIONS

Conceptualization: DP, CC, LPdA; Investigation: DP, JC-S, AV-F, JD-N, DDL, MIM; Methodology: DP, IO, VC, CAA, SML, NG; Writing, original draft: DP; Reviewing, and editing: LPdA, CC, SML, JD-N, NG, DDL, MIM, AV-F, CAA.

## ACKNOWLEDGMENTS

We would like to thank Luisa Cortes for all the help in the nucleus circularity measurements – copyright © 2015 Jorge Valero Gomez-Lobo & Luísa Cortes - MICC Imaging facility of CNC, including Margarida Caldeira, Tatiana Catarino (for assistance with microscopy imaging). We also would like to acknowledge David Rufino-Ramos, Magda M Santana and all members of the LPdA lab for all the support, discussions, and comments.

Schematic figures were partially created using <u>Biorender.com</u>

This work was funded by the European Commission Seventh Framework Programme FP7/2010 under Grant 264508 (TreatPolyQ) and the Federal Ministry of Education and Research (PPPT-MJD Grant 01GM1309B) under the E-Rare-2 [ERA (European Research Area) - Net for research programmes on rare diseases, ERDF through the Regional Operational Program Center 2020, Competitiveness Factors Operational Program (COMPETE 2020), and national funds through FCT (Foundation for Science and Technology) - BrainHealth2020 projects (CENTRO-01-0145-FEDER-000008), UID/NEU/04539/2020, LA/P/0058/2020, ViraVector (CENTRO-01-0145-FEDER-022095), CortaCAGs (PTDC/ NEU-NMC/0084/2014|POCI-01-0145-FEDER-016719), SpreadSilencing POCI-01-0145-FEDER-029716, CENTRO-01-0246-FEDER-000010 (MIA-Portugal), Imagene POCI-01-0145-FEDER-016807, CancelStem POCI-01-0145-FEDER-016390 and POCI-01-0145-FEDER-032309, as well as SynSpread, ESMI, and ModelPolyQ under the EU Joint Program-Neurodegenerative Disease Research (JPND), with the last two co-funded by the European Union H2020 program, GA no. 643417 and no. 857524; E-Rare4/0003/2012 Joint Call for European Research Projects on Rare Diseases (JTC 2012) and FCT; by the National Ataxia Foundation (USA), the American Portuguese Biomedical Research Fund (APBRF), and the Richard Chin and Lily Lock Machado-Joseph Disease Research Fund. DP was supported by a PhD fellowship from FCT (SFRH / BD / 51965 /2012); AV-F – (SFRH/BD/87804/2012); SML – (SFRH/BD/51673/2011); MIM – (PD/BD/114171/2016); DDL-(2020.09668.BD).

## DATA AND MATERIALS AVAILABILITY

All data needed to evaluate the conclusions in the paper are present in the paper and/or the Supporting information. Additional data related to this paper may be requested from the authors.

