## Supporting information for "Nuclear aging in polyglutamine-induced neurodegeneration"

#### Extended Experimental Procedures

##### 4.5 Western Blot

After incubating with the primary antibodies, the membranes were then incubated with the corresponding Alkaline Phosphatase conjugated secondary antibodies: goat anti-mouse (1:10,000; #31328, Thermo Scientific Pierce), rabbit anti-goat (1:10,000; sc2771 Santa Cruz Biotechnology) and goat anti-rabbit (1:10,000; Thermo Fisher Scientific) for 2 hours at RT. Bands were visualized with Enhanced Chemifluorescent substrate (ECF) (GE Healthcare), using chemifluorescence imaging (Chemidoc imaging system, Biorad). Semi quantitative analysis was carried out based on the optical density (OD) of scanned films (Image lab software version 5.1; Biorad, CA, USA). The specific optical density was then normalized with respect to the amount of β-actin/β-tubulin loaded in the corresponding lane of the same gel.

##### 4.6 Histological staining and immunohistochemical analysis

###### Bright-field Immunohistochemistry

After incubation with primary and secondary antibodies, bound antibodies were visualized using the VECTASTAIN ABC kit (PK-6100, Vector Laboratories), with 3',3'-diaminobenzidine tetrahydrochloride (DAB) as substrate (SK-4100, Vector Laboratories). Sections were then placed in gelatine-coated slides, dried and dehydrated and cleared with xylene, and mounted. Staining was visualized, and images captured with the Zeiss Axioskop 2 imaging microscope (Carl Zeiss MicroImaging microscope, Carl Zeiss MicroImaging) equipped with AxioCam HR colour digital cameras (Carl Zeiss Microimaging) and Plan-Neofluar 5x/0.15 Ph 1 (440321), Plan-Neofluar 20x/0.50 Ph 2 (1004-989), Plan-Neofluar 40x/0.75 Ph 2 (440351) and Plan-Neofluar 63x/1.25 Oil (440460-0000) objectives using the AxioVision 4.7 software package (Carl Zeiss Microimaging).

#### 4.7 Immunofluorescent analysis

##### Fluorescent immunocytochemistry

After incubating with the primary antibodies, and washing with PBS, cells were incubated with donkey anti-goat Alexa Fluor 488 and donkey anti-rabbit, or mouse Alexa Fluor 555 secondary antibodies (1:250, Invitrogen) for 2 hours at RT. Additionally, cells were stained with 4′,6-Diamidine-2′-phenylindole dihydrochloride (DAPI, 1:5000, Sigma-Aldrich) and after washing, mounted in Fluoroshield (Sigma-Aldrich). Fluorescent staining was detected using a Zeiss motorized inverted widefield microscope (Carl Zeiss Axio Observer Z1), equipped with a CCD digital camera (Axiocam HRm) and a digital CMOS camera (ORCA Flash 4.0) and with a Plan-Apochromat 63x/1.40 Oil DIC (UV) M27 objective. Images were acquired and processed using the ZEN 2 Blue edition software (Carl Zeiss Microscopy GmbH, 2011)

##### Fluorescence Immunohistochemistry

After incubation with primary antibodies, sections were washed and incubated for 2 hours at RT with the corresponding secondary antibodies coupled to fluorophores: goat anti-mouse Alexa Fluor 647 or Alexa Fluor 594, goat anti-rabbit Alexa Fluor 568 (1:250, Molecular Probes-Invitrogen, Eugene, OR), diluted in blocking solution. The sections were then washed and incubated for 10 minutes with 4′,6-Diamidine-2′-phenylindole dihydrochloride (DAPI; 1:5000, Sigma-Aldrich, St. Louis, USA), washed, and mounted in fluorescence mounting medium (Dako) on microscope slides.

Staining was visualized, and images captured with the Carl Zeiss Axio Imager Z2 (Carl Zeiss MicroImaging, Oberkochen, Germany), equipped with a CCD monochromatic digital camera (Axiocam HRm) and an EC Plan-Neofluar 10x/0.30M27 objective, using the ZEN 2 Blue edition software (Carl Zeiss Microscopy GmbH, 2011).

##### Fluorescence Immunohistochemistry - yH2AX

After 36h incubation with primary antibody, sections were washed and incubated for 2 hours at RT with the secondary antibody goat anti-mouse 594 (1:250, Molecular Probes-Invitrogen, Eugene, OR) diluted in the same blocking solution as the primary antibody. Sections were washed, and nuclei were counterstained with DAPI (1:5000) for 10 minutes at RT, washed, and mounted with Fluorescence Mounting Medium (S302380-2, Dako). Staining was visualized with Carl Zeiss Axio Imager Z2 (Carl Zeiss MicroImaging, Oberkochen, Germany), equipped with a CCD monochromatic digital camera (Axiocam HRm) and a CCD colour digital camera (Axiocam HRc), and using the ZEN 2 Blue edition software (Carl Zeiss Microscopy GmbH, 2011).

#### 4.8 Imaging Analysis

##### Nuclear circularity assessment

Nuclear circularity was analysed using Fiji (Fiji Is Just ImageJ, NIH) - Measure nucleus single channel image – copyright © 2015 Jorge Valero Gomez-Lobo and Luísa Cortes-MICC Imaging facility of CNC.

##### Quantification of pyknotic nuclei

After cresyl-violet staining, the number of pyknotic nuclei in the area under the needle tract was counted. Six specific regions under the needle tract of each injection were photographed with a 40x objective. The total number of the pyknotic nuclei in this area was blindly counted for each animal and each hemisphere, using a semiautomated image analysis software Fiji (Fiji Is Just ImageJ, NIH).

##### Quantification of mutant ataxin-3 ubiquitinated inclusions

Twelve coronal sections (with 200 µm intervals from each other) and showing the entire striatum were scanned with a 20x objective. These sections covered the entire region containing mutant ataxin-3 ubiquitinated inclusions, as revealed by the staining with an anti-ubiquitin antibody. The total number of ubiquitinated inclusions was estimated by the formula: total number = s (n1+n2+n3+…+n12), where “s” represents the number of intermediate sections (8), and “n1-n12” represents the number of inclusions present in each section. All inclusions were manually counted using a semiautomated image analysis software package Fiji (Fiji Is Just ImageJ, NIH) and ZEN 2 Blue edition software (Carl Zeiss Microscopy GmbH, 2011).

##### Quantification of DARPP-32 depleted volume

The extent of striatal DARPP-32 loss was made by screening 12 stained coronal sections per animal (distanced 200 µm from each other) with a 5x objective and selected in order to obtain a sampling of all striatum. The area of the lesion was quantified with a semi-automated image-analysis software package Fiji (Fiji Is Just ImageJ, NIH). The area of the striatum showing a loss of DARPP-32 staining was measured for each animal, with an operator-independent macro. The volume was then estimated with the following formula: volume = d (a1+a2+…+a11+a12), where “d” is the distance between serial sections (200 µm) and “a1-a12” are DARPP-32 depleted areas for individual serial sections, as previously described (de Almeida et al., 2002). The depleted area corresponds to the area with a gray-scale value lower than the mean gray-scale value of all pixels measured around the lesioned area.

##### Quantification of GFAP (astrogliosis) area and AIF1/Iba-1 immunoreactivity (microglia recruitment)

Protein's immunoreactivity quantification and analysis were made as previously described (Gonçalves et al., 2013).

Quantitative analysis of fluorescence was performed with ZEN 2 Blue edition software (Carl Zeiss Microscopy GmbH, 2011) and images were taken under identical image acquisition conditions. Uniform adjustments of brightness and contrast were made to all images.

##### Quantification of microglial recruitment (number of cells)

Quantification of microglial recruitment (number of AIF1/Iba-1-positive cells) was quantified with a semi-automated image-analysis software package Fiji (Fiji Is Just ImageJ, NIH). After manually thresholding images to detect AIF1/Iba-1-positive cells (by applying the "watershedding" tool), particles ranging 10-500 $\mu m^2$ were automatically selected and counted (scale: 2 pixels/µm). The estimative calculation of the total number of cells in the entire striatum was performed as previously described for mutant Atxn3 ubiquitinated inclusions.

##### Quantification of yH2AX foci - DNA damage marker

Coronal sections showing the entire striatum (12 sections per animal) were scanned with a 20x objective. The analysed areas of the striatum stained with an antibody against γH2AX, covered the entire transduced region. All foci were automatically analysed and counted using the software package Fiji (Fiji Is Just ImageJ, NIH) and ZEN 2 Blue edition software (Carl Zeiss Microscopy GmbH, 2011), by manually thresholding images. Particles ranging 2-400 $\mu m^2$ were automatically analysed and counted (scale: 2 pixels/µm). The total number of striatal γH2AX foci was estimated by using the same formula as previously described for mutant Atxn3 ubiquitinated inclusions.

### Supporting Figures

#### Figure S1

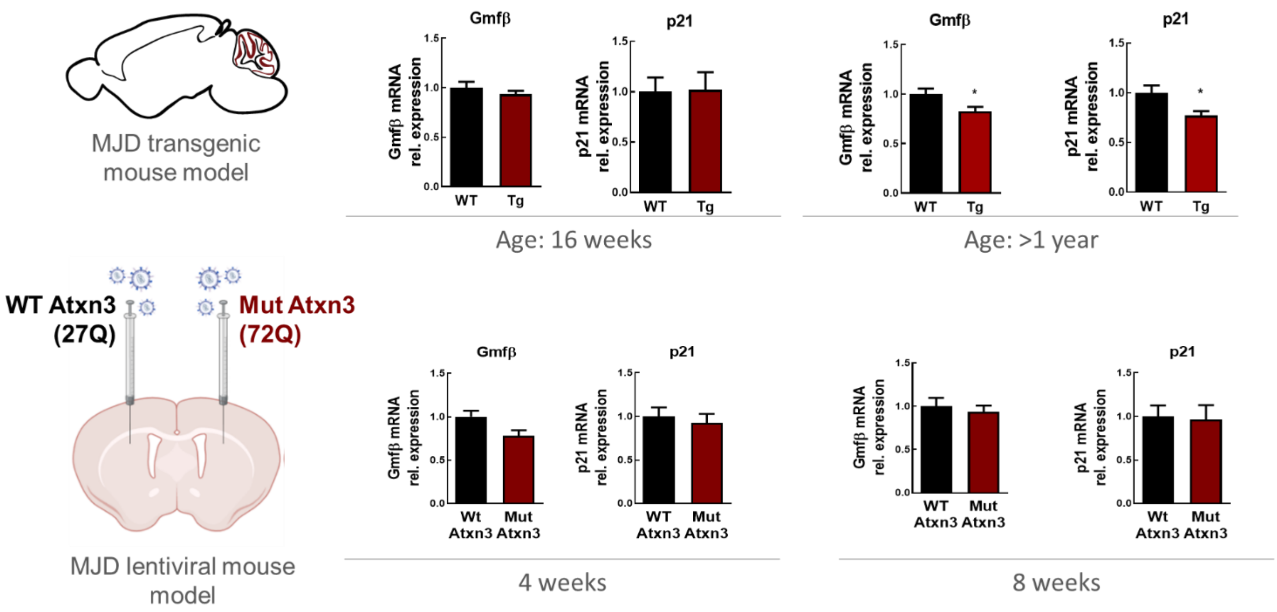


Figure S1. **Expression of Gmfβ and p21 in MJD models**. Gmfβ and p21 mRNA levels evaluated by qRT-PCR in **(a)** MJD transgenic mouse model and **(f)** MJD lentiviral-based mouse model, submitted to the same procedures described in Figure 1. Significant decrease of Gmfβ and p21 mRNA levels **(d, e)** in the cerebellum of MJD transgenic mice with > 1 year of age. Transgenic mice with 16 weeks of age and the MJD lentiviral mouse model did not show any alterations (**b, c, g-j**) qRT-PCR analysis was normalized with endogenous control (Gapdh). Statistical significance was evaluated with Unpaired t-test *$p<0.05$, n=5 (Tg - 16 weeks); n=7/5 (Tg>1y); and paired Student's t-test n = 5 (LV model). Data are expressed as mean ± SEM. Gmfβ - Glial maturation factor β.

**Figure S2**

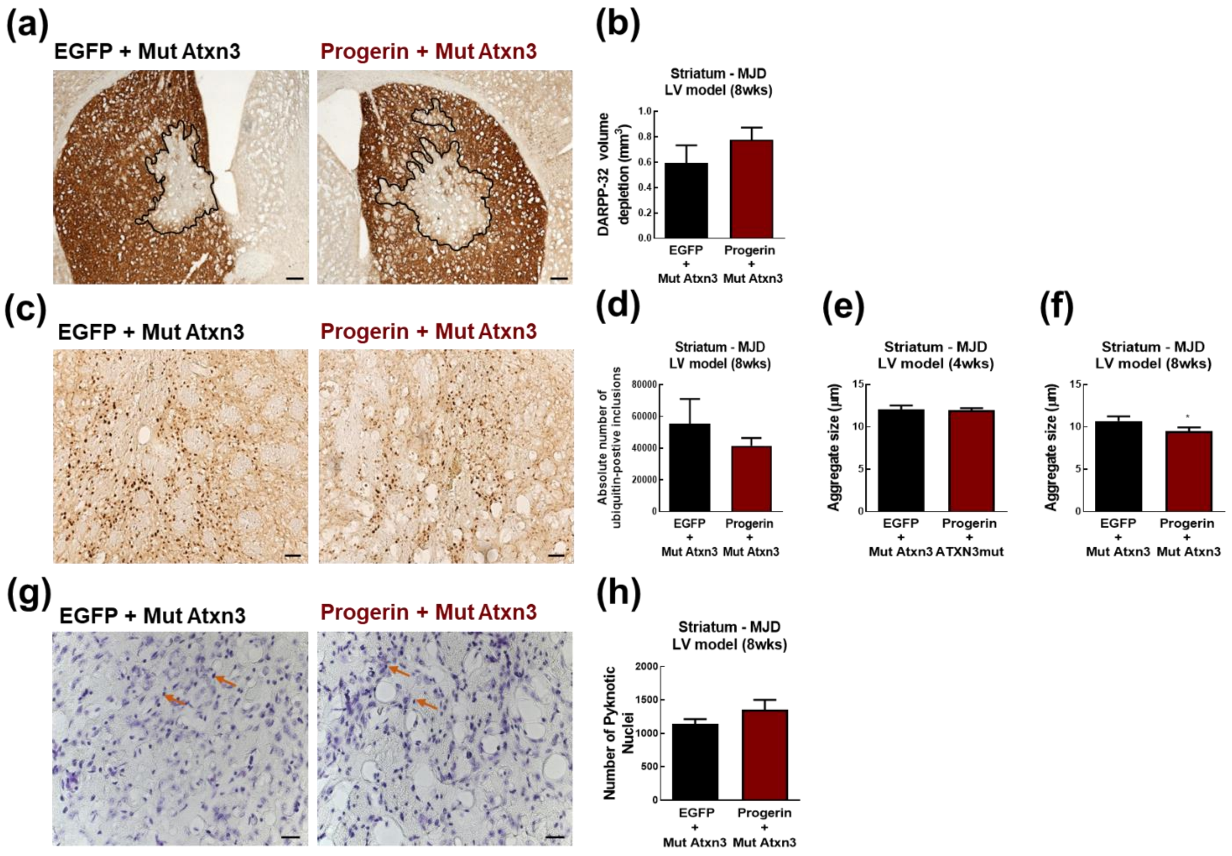


Figure S2. **Progerin overexpression and neuropathology in a lentiviral vector-based mouse model of MJD, 8 weeks post-injection**. **(a)** Immunohistochemical peroxidase staining using anti-DARPP32 antibody as dopaminergic loss of function marker, and depleted volume quantification **(b)**. No significant alterations in DARPP-32 volume depletion among hemispheres, 8-week post-injection. **(c)** Immunohistochemical analysis using anti-ubiquitin antibody and quantification of positive-ubiquitin inclusions **(d)** with progerin injected hemisphere displaying a significantly decrease in aggregate size, 8 weeks post-injection **(f)** and no differences found, 4 weeks post-injection **(e)**. **(g)** Cresyl violet staining indicating pyknotic nuclei (orange arrows), and its quantification **(h)**. Statistical significance was evaluated with paired Student's t-test, n=5. Data are expressed as mean ± SEM. Scale bars represent **(a)** 500µm, **(c)** 100µm, and **(g)** 200µm.

**Figure S3**

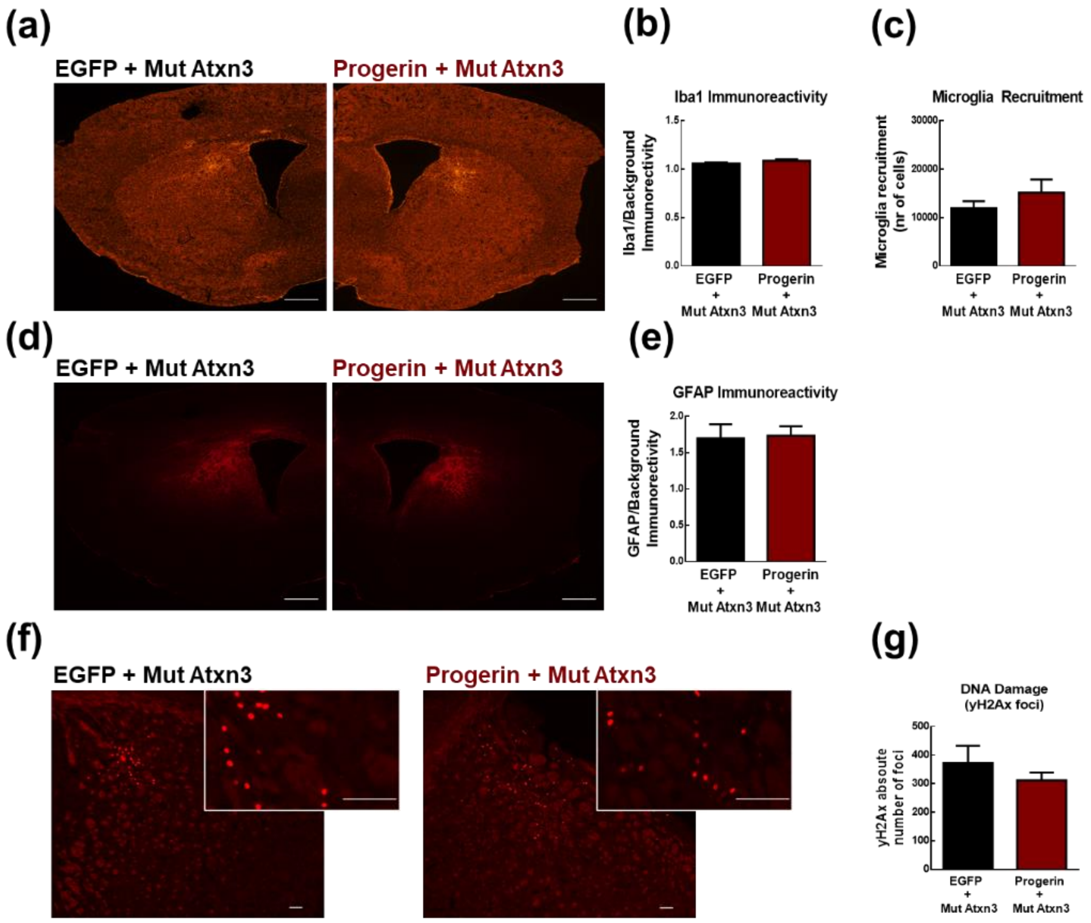


**Figure S3 - Progerin overexpression and its role in neuroinflammation and in DNA damage, 8 weeks post-injection.** Mice were submitted to the same procedure described in Figure 4. **(a)** Fluorescent immunohistochemical analysis of Iba-1, **(b)** relative fluorescence immunoreactivity measurement among hemispheres and **(c)** microglial recruitment, with no detectable alterations at 8 weeks post-injection. **(d)** Fluorescent immunohistochemical analysis of GFAP in the striatum, showing no alterations among hemispheres **(e)**. **(f)** Fluorescent immunohistochemistry for the detection and quantification of DNA damage using the phosphorylation of γH2AX as a marker/antibody, presenting no alterations among hemispheres **(g)**. Microglial recruitment and γH2AX foci measured using Fiji (Fiji Is Just ImageJ, NIH). Statistical significance was evaluated with paired Student's t-test, n=5. Data are expressed as mean ± SEM. Scale bars represent **(a,d)** 500µm, **(f)** 100µm.

**Figure S4**

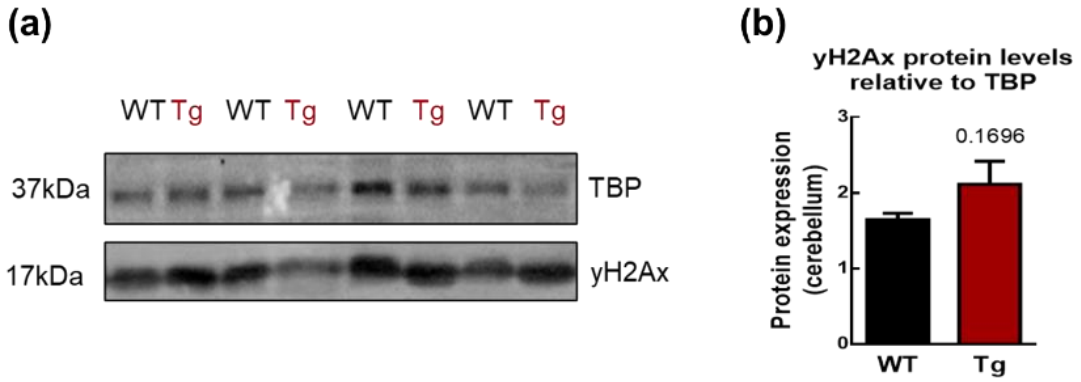


Figure S4. **DNA damage in transgenic MJD model**. yH2Ax protein levels in MJD transgenic mouse model quantified through WB. Tissues of MJD transgenic mice with 16 weeks were collected and processed by fractioning protocol for WB **(a)**. All protein relative levels were normalized with TBP **(b)**. Statistical significance was evaluated with paired Student's t-test, n = 6. Data are expressed as mean ± SEM. TBP – TATA binding Protein.
